# *Mmp17-*deficient mice exhibit heightened goblet cell effector expression in the colon and increased resistance to chronic *Trichuris muris* infection

**DOI:** 10.1101/2023.07.10.548379

**Authors:** Pia M. Vornewald, Ruth Forman, Rouan Yao, Naveen Pamar, Håvard T. Lindholm, Mara Martín-Alonso, Kathryn J. Else, Menno J. Oudhoff

## Abstract

Intestinal epithelial homeostasis is maintained by intrinsic and extrinsic signals. The extrinsic signals include those provided by mesenchymal cell populations that surround intestinal crypts and is further facilitated by the extracellular matrix (ECM), which is modulated by proteases such as matrix metalloproteinases (MMPs). Extrinsic signals ensure an appropriate balance between intestinal epithelial proliferation and differentiation. This study explores the role of MMP17, which is expressed by mesenchymal cells, in intestinal homeostasis and during immunity to infection. Mice lacking MMP17 expressed high levels of goblet-cell associated genes, such as CLCA1 and RELM-β, which are normally associated with immune responses to infection. Nevertheless, *Mmp17* KO mice did not have altered resistance during a bacterial *Citrobacter rodentium* infection. However, when challenged with a low dose of the helminth *Trichuris muris*, *Mmp17* KO mice had increased resistance, without a clear role for an altered immune response during infection. Mechanistically, we did not find changes in traditional modulators of goblet cell effectors such as the NOTCH pathway or specific cytokines. Instead, we found elevated BMP signaling in *Mmp17* KO mouse large intestinal crypts that we propose to alter the goblet cell maturation state. Together, our data suggests that MMP17 extrinsically alters the goblet cell maturation state *via* a BMP signaling axis, which is sufficient to alter clearance in a helminth infection model.

## INTRODUCTION

The intestinal tract is lined by a single layer of intestinal epithelial cells. This intestinal epithelium comprises multiple cell types, each with distinct functions, that work together to provide a barrier against harmful microorganisms, while at the same time allowing for nutrient and water absorption.(1) To fulfill those functions the intestinal epithelium is constantly replenished by a pool of LGR5+ stem cells located at the base of epithelial invaginations known as intestinal crypts, with a turnover rate of 3-5 days.(2, 3) This remarkable regenerative capacity enables the intestinal epithelium to rapidly respond to injury or infection.

Goblet cells play a critical role in maintaining the intestinal homeostasis by producing key components of the mucus layer, such as MUC2, CLCA1, and FCGBP.(4-7) The mucus serves as a physical barrier, preventing the attachment and penetration of bacteria(8) and other pathogens, including helminths.(9) Additionally, the mucus contains proteins with antimicrobial properties like RELM-β and ANG4, which further bolsters the gut’s innate immune defense.(10-14) In response to infections both the amount of mucus and its composition can be altered, with goblet cell specific antimicrobial proteins (AMPs) increasing due to immune-cell derived cytokines that directly induce signaling in intestinal epithelium.(15) In fact, an increase in goblet cell numbers is crucial in the defense against pathogenic helminths.(9)

Epithelial homeostasis depends on a niche created by gradients of factors to preserve stemness at the base of crypts while enabling the differentiation of mature and functionally diverse cell types towards the lumen. Critical niche factors include WNTs, R-spondins, Bone morphogenic proteins (BMPs), and prostaglandins, which are expressed by mesenchymal cell subtypes and immune cells such as macrophages that are present in the lamina propria between the epithelium and the smooth muscle layers.(16-22)

In addition to secreted factors, the structure and composition of the extracellular matrix (ECM) within the lamina propria also regulates the intestinal niche.(23) Matrix metalloproteinases (MMPs) are a key group of ECM regulators, modifying the structure and function of the ECM both through direct action on ECM components and by cleaving ECM associated-molecules, such as growth factors.(24) MMPs have been implicated in wound healing, (chronic) inflammation, and cancer.(25-27) We recently demonstrated that the membrane-bound MMP, MMP17 (also called MT4-MMP) is important in maintaining epithelial homeostasis and its ability to respond to injury. Furthermore, we identified the intestinal smooth muscle as a source of niche factors.(28)

In the present work, we show that the loss of *Mmp17* induces goblet cell effector genes in the colon and cecum epithelium. We propose that these are caused by elevated BMP signaling in the epithelium and that these changes to the epithelium result in improved clearance of low-dose *Trichuris muris* infection.

## RESULTS

### Mmp17 KO mice show increased expression of goblet cell-effectors in proximal colon goblet cells

We recently demonstrated that mice lacking the membrane-bound matrix metalloproteinase MMP17 display impaired intestinal regenerative capacities in both a chemically-induced colitis and whole-body irradiation injury models. In our previous study, we conducted transcriptomic analysis of colon crypts isolated from naïve *Mmp17+/+* (WT) and *Mmp17-/-* (KO) mice using bulk RNA sequencing (RNA-seq).(28) We re-examined this dataset and identified several goblet cell-associated genes among the most upregulated genes in crypts derived from KO animals, particularly *Ang4, Clca1, Retnlb*, and *Muc2* (Figure 1a). *Retnlb, Clca1*, and *Ang4* can be classified as AMPs, which are typically upregulated in the intestine during infections. Immunofluorescence (IF) staining of RELM-β, the protein encoded by the *Retnlb* gene, confirmed increased protein levels and revealed a distinct pattern of RELM-β positive cells in the proximal region of the colon (Figure 1b). Similarly, we observed a higher fluorescent intensity in KO tissue compared to WT tissue in the proximal colon when stained for CLCA1 (Figure 1c). Examination of other AMP genes, including *Sprr2a1, Sprr2a2*, and *Sprr2a3*, showed slightly increased levels, albeit not significant (Supplementary Figure 1a).(29) RNA levels of the core mucus protein *Fcgbp* were also significantly elevated in KO epithelium (Supplementary Figure 1b). The upregulation in goblet cell-associated genes in KO animals did not correlate with an increased number of this cell type, as demonstrated by the quantification of total goblet cell numbers per crypt in WT and KO mouse colon using alcian blue staining (Figure 1d). Accordingly, we detected no changes in the expression of the transcription factors *Atoh1* and *Spdef* (Figure 1e), which are involved in NOTCH signaling to regulate goblet cell differentiation. Since our previous data indicated that MMP17 is not expressed in the intestinal epithelium, we sought to determine whether the increase in epithelial goblet cell genes was intrinsic to the epithelium or due to changes in the surrounding niche *in vivo*. We isolated colonic crypts from WT and KO mice to cultivate colon organoids (colonoids), measured *Retnlb* expression via qPCR (Figure 1f) and assessed RELM-β protein expression using IF staining (Figure 1g). We observed no differences in RELM-β protein and *Retnlb* mRNA expression, suggesting that the increased expression of goblet cell-associated genes in KO mouse crypts is not intrinsic to the intestinal epithelium but rather a consequence of the environment in the surrounding niche. In summary, we find that animals lacking MMP17 exhibit elevated levels of goblet cell effector proteins without an increase in goblet cell numbers, that are presumably induced by extrinsic factors in the proximal colon.

**Figure 1:**
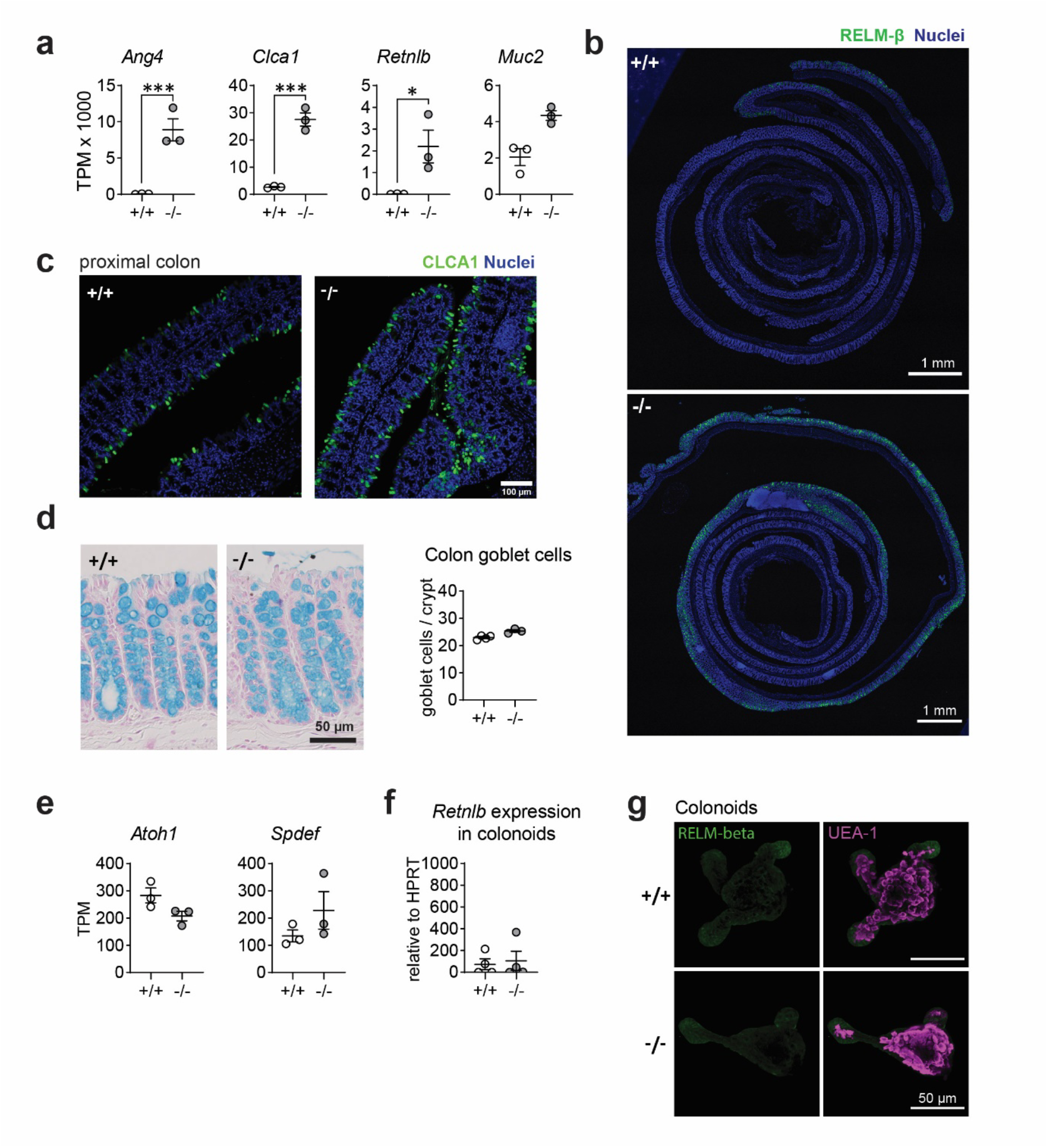
MMP17 controls goblet cell expression in colon. (**a**) Graphs of RNAseq data of naive mouse colon crypts from WT (*+/+*) and *Mmp17* KO (*-/-*) mice depicting the expression levels of *Ang4*, *Clca1, Retnlb* and *Muc2*, n=3, padj calculated with DESeq2, * p<0.05, ** p<0.01, *** p<0.001. (**b**) Immunofluorescence (IF) images of Colon Swiss rolls of naive mouse full-length colon of WT (*+/+*) and *Mmp17* KO (*-/-*) mice, for RELM-β (green), nuclear stain (blue), proximal end on the outside, scale 1 mm. (**c**) IF stain images of naive mouse proximal colon of WT (*+/+*) and KO (*-/-*) mice for CLCA1 (green), nuclear stain (blue), scale 100 µm. (**d**) The pictures on the left show Alcian blue stain for goblet cells, on the right is a count of goblets/crypt in proximal colon of naive WT (*+/+*) (n=4) and KO (*-/-*) (n=3) mice, 30 crypts counted for each animal. (**e**) Graphs of RNAseq data of naive mouse colon crypts from Mmp17 +/+ and -/- mice showing the expression for *Atoh1* and *Spdef*, n=3, padj * p<0.05, ** p<0.01, *** p<0.001. (**f**) Graph of qPCR data for *Retnlb* of WT (*+/+*) and KO (*-/-*) mouse colonoids after one split, Ct value relative to *HPRT,* n=4. (**g**) IF staining images for RELM-β in WT (*+/+*) and KO (*-/-*) mouse colonoids after one split, RELM-β (green) and UEA-1 (magenta), scale 50 µm. Numerical data are means ± SEM. Data in (a) and (e) represents p-adjusted value from RNAseq analysis, and data in (d) and (f) was analyzed using Student’s unpaired t-test (two-tailed).

### Goblet cell effector upregulation is not caused by a change in cytokine levels

*Ang4*, *Clca1,* and *Retnlb* are goblet-cell associated genes that are induced during immune responses, such as bacterial and helminth infections, and are typically stimulated by type 2 and 3 immune cytokines such as IL-4/IL-13 (type 2) and IL-22 (type 3).(10, 11, 14, 15, 30) To investigate whether naïve KO animals inherently exhibit elevated cytokine levels, we measured levels of IL-13, IL-22, and IFN-γ in re-stimulated mesenteric lymph node lymphocytes (mLNL) *in vitro* and homogenized colon tissue using ELISA (Figure 2a). We also isolated mRNA from whole proximal colon tissue and measured gene expression of the same cytokines (Figure 2b). Our analysis revealed no significant differences in the levels of IL-13 and IL-22 protein (mLNL, colon) or gene expression (colon) between wild-type (WT) and knockout (KO) mice. Of note, in all cases, we did observe a subtle increase in the levels of IFN-γ/*Ifng*, but only with a significant elevation in colon tissue (Figure 2a,b). While IFN-γ is not known to induce goblet cell effectors or their differentiation in intestinal epithelium (15), it has been reported that RELM-β can stimulate IFN-γ production in T-cells. (31) Therefore, we propose that the modest increase in IFN-γ levels may be a result of RELM-β upregulation rather than the inverse. To further rule out the possibility that the observed increase in goblet cell effector proteins was due to IL-4/IL-13 or IL-22 signaling, which are known to induce such effectors, we also analyzed the levels of phosphorylated (p)STAT6 and pSTAT3, which are downstream signaling molecules of IL-4/IL-13 signaling and IL-22 signaling respectively. We observed no significant difference in protein levels of pSTAT6 (Figure 2c) or pSTAT3 (Figure 2d) between WT and KO colon crypts, thus excluding IL-4/IL-13 and IL-22 signaling as the cause for the altered epithelial expression. In conclusion, the heighted levels of goblet cell effectors in Mmp17 KO mice are not the result of a heightened level of cytokines during homeostasis.

**Figure 2:**
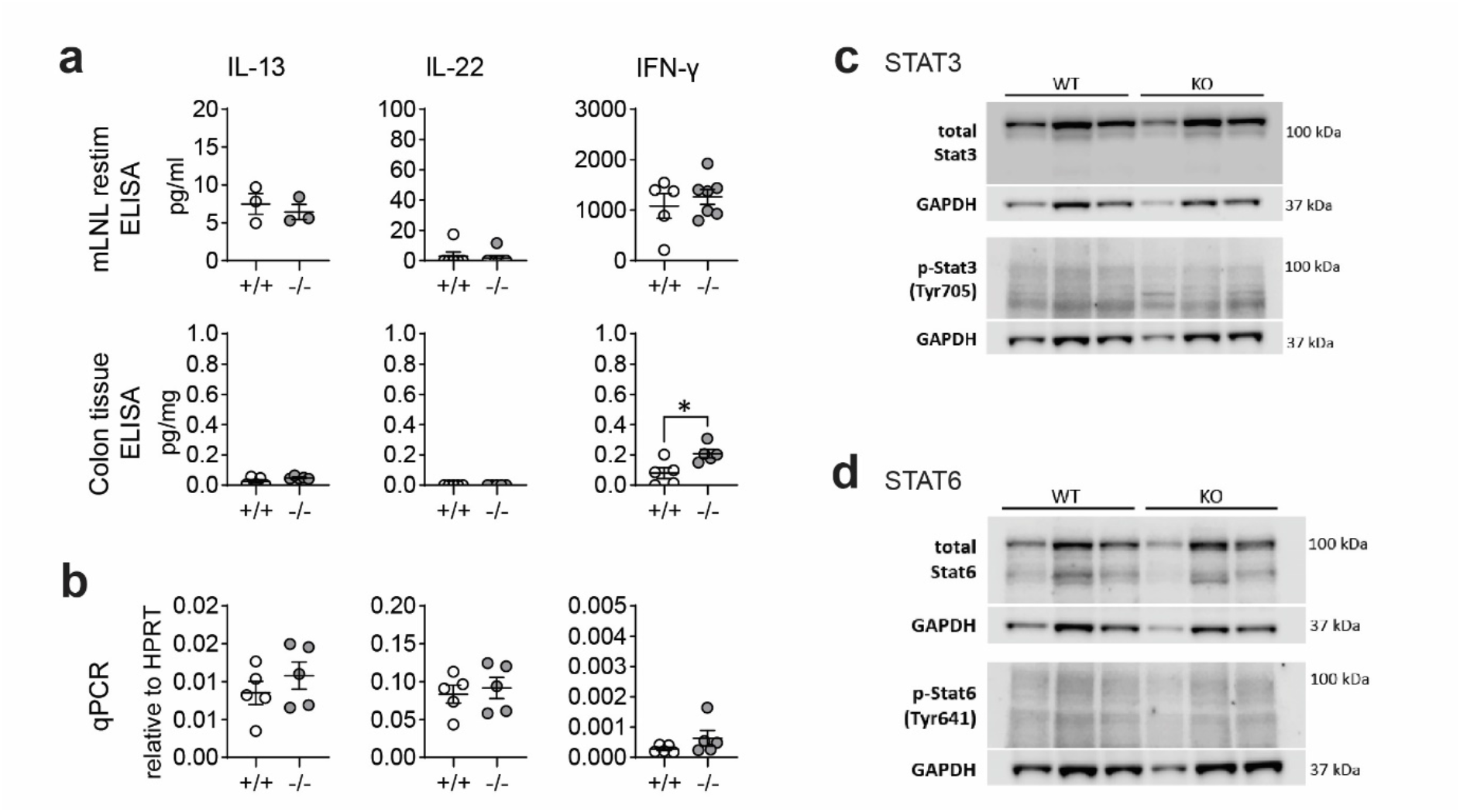
*Mmp17* KO mice do not have increased cytokine levels in the colon. (**a**) Graphs show cyokine ELISA of mLNLs restimulations (first row) for IL-13, IL-22 and IFNγ, cytokine ELISA of proximal colon tissue adjusted to mg tissue (second row) for IL-13, IL-22, and IFNγ. **(b)** qPCR for *Il13, il22* and *Ifng* in proximal colon tissue relative to *Hprt* (third row), n=3-6. **(c)** Western blots of protein isolates from colon crypts for STAT3 and pSTAT3, GAPDH loading control, n=3. (**d**) Western blots of protein isolates from colon crypts for STAT6 and pSTAT6, GAPDH loading control, n=3. Numerical data are means ± SEM. Data were analyzed using Student’s unpaired t-test (two-tailed). * p<0.05

### Increased expression of goblet cell effector proteins does not influence the course of *Citrobacter rodentium* infection

Goblet cell effector proteins, which are secreted into the intestinal lumen as part of the mucus layer, form a crucial line of defense against pathogens in the intestine. We hypothesized that the elevated levels of these effectors might offer an advantage in clearing intestinal infections, as both ANG4 and RELM-β have been described to have anti-bacterial properties.(11, 32) Consequently, we infected WT and KO mice with *Citrobacter rodentium*, a gram-negative bacterial pathogen that primarily infects the colon, causing mild colitis and diarrhea. Generally, mice with a C57BL/6 background exhibit resistance to *C. rodentium* and can clear the pathogen in 2 to 4 weeks, depending on the infectious dose.(33, 34) Both WT and KO-infected mice cleared the infection between 11-20 days. Although the average duration for clearance was shorter in KO mice (14.7 dpi) compared to WT mice (16.7 dpi), these differences were not significant (Figure 3a). Furthermore, we observed equal peak bacterial burdens at day 12 post-infection, averaging at 10^8^ colony-forming units (CFU) per gram (g) of fecal matter for all mice (Figure 3b). The immune response to *C. rodentium* is characterized by the induction of IL-22 and IL-17A which contribute to the clearance.(35) We noted similar levels of IL-22 and IL-17A in both WT and KO animals at day 12 indicating there was no difference in mounting a type 3 response (Figure 3c). Crypt hyperplasia, resulting from increased epithelial proliferation is a hallmark of *C. rodentium* infection. As we previously discovered that epithelial proliferation was nearly absent in KO animals after chemical-induced colitis-type injury in the colon(28), we measured the crypt length and Ki67 staining to assess proliferation. We observed no changes in either crypt length or Ki67+ staining (Figure 3d and 3e). In conclusion, despite KO animals displaying heightened homeostatic levels of goblet cell effectors, including bacteria-targeting AMPs like RELM-β (11, 29), this does not impact immunity to *C. rodentium* infection.

**Figure 3:**
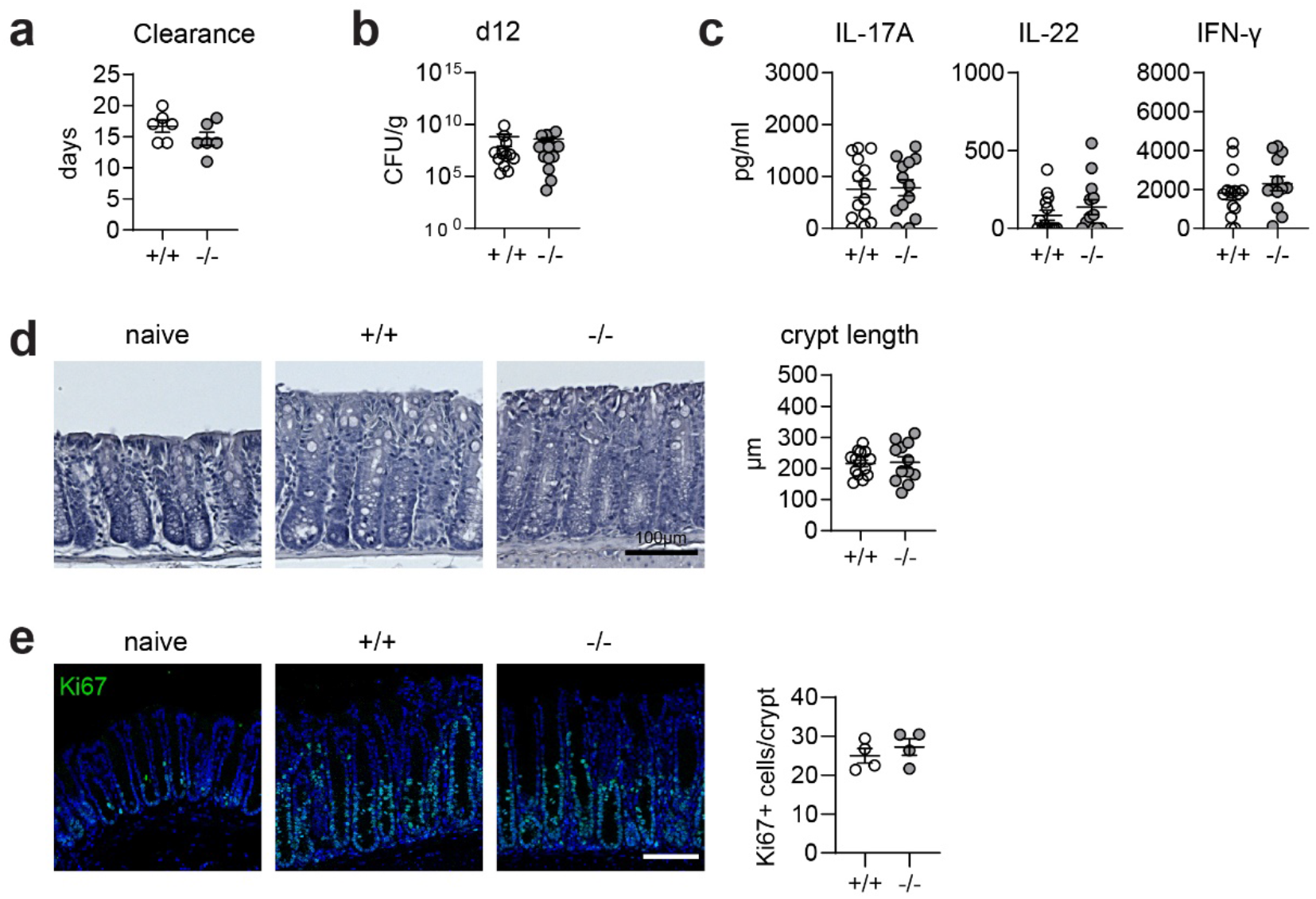
*C. rodentium* infection is not altered in *Mmp17* KO mice. (**a**) Graph shows days until clearance of *C. rodentium* measured using fecal CFUs, single experiment, n=6. (**b**) Quantification of fecal CFUs at 12 days post infection (dpi) with *C. rodentium* is depicted in the graph, 2 independent experiments pooled, +/+ (n=14), -/- (n=13). (**c**) Cyokine levels measured via ELISA of mLNLs restimulations for IL-13, IL-22, and IFNg. (**d**) Images depicting hematoxylin and eosin (H&E) staining of naive and *C. rodentium* WT (*+/+*) and KO (*-/-*) mouse distal colon 12 dpi, scale 100 µm (left) and crypt length quantification in µm (right), 2 independent experiments pooled, +/+ (n=14), -/- (n=13). (**e**) Confocal maximal projection images showing Ki67 IF stain of naive and *C. rodentium* infected WT (*+/+*) and KO (*-/-*) mouse distal colon 12 dpi, scale 100 µm, Ki67 (green), nuclear stain (blue). Numerical data are means ± SEM. Data was analyzed using Student’s unpaired t-test (two-tailed).

### Increased expression of goblet cell effector proteins does not influence the course of a high dose *Trichuris muris* infection

Clearance of parasitic worms (helminths) in the gut relies on a “weep and sweep” response induced by type 2 cytokines (IL-4, IL-5, IL-13), which includes increased epithelial turnover, upregulated mucus production by goblet cells, and enhanced muscle contraction, collectively working to expel the parasites.(36) Goblet cell effectors CLCA1, ANG4, and RELM-β are induced during infection with the helminth *Trichuris muris*.(9, 30, 37, 38) CLCA1 plays a role in the formation of the two distinct mucus layers in the colon and is upregulated upon *T. muris* infection(39), while RELM-β can reduce the viability of *T. muris*(10) and ANG4 expression has been correlated with the expulsion of *T. muris*.(14, 40) Given heightened levels of these effector proteins during homeostasis, we hypothesized that KO animals would be better equipped to combat *T. muris* infection compared to WT animals.

Male mice on a C57BL/6 background exhibit a dose-dependent immune response to *T. muris*. Depending on the infective dose of *T. muris,* the immune system is polarized towards a susceptibility-associated type 1 immune response (low dose) or a resistance-associated type 2 immune response (high dose)(36). Initially, we infected both WT and KO male mice with a high dose of *T. muris* eggs. After 14 days all mice developed an infection, which was mostly cleared after 21 days (Figure 4a). During *T. muris* infection, increased epithelial proliferation leads to crypt hyperplasia. Histological analysis showed comparable crypt length at both day 14 and day 21 (Figure 4b), and Ki67 staining indicated equivalent increased proliferation at d21 (Figure 4c) in both WT and KO animals. As observed during *C. rodentium* infection, the previously described lack of regenerative ability of the KO epithelium does not impair increased epithelial proliferation during infection. Counts of RELM-β+ cells were not significantly different d21 days post infection (Figure 4d). To assess the immune response, we stimulated mLN with E/S antigen and measured the cytokine levels for IFNg, IL-6, IL-10, IL-4, IL-5, and IL-13 (Figure 4e). Increased levels of IL-13 and IL-5 in re-stimulated mLN cells as well as serum IgE indicated a typical and equal type 2 immune response in both WT and KO animals (Figure 4e and 4f). Mice with a type 2 cytokine profile produce high levels of IgG1, while mice with increased levels of type 1 cytokines exhibit elevated levels of IgG2. Although there was no difference in IgG1 levels, *Mmp17* KO mice showed significantly higher, through variable, levels of IgG2 despite having comparable type 1 cytokine levels to WT animals (Fig 4g). In conclusion, while the expression of AMPs and mucus proteins during *T. muris* infection has been shown to be essential(29), the higher baseline levels in *Mmp17* KO mice do not influence the course of high dose *T. muris* infection.

**Figure 4:**
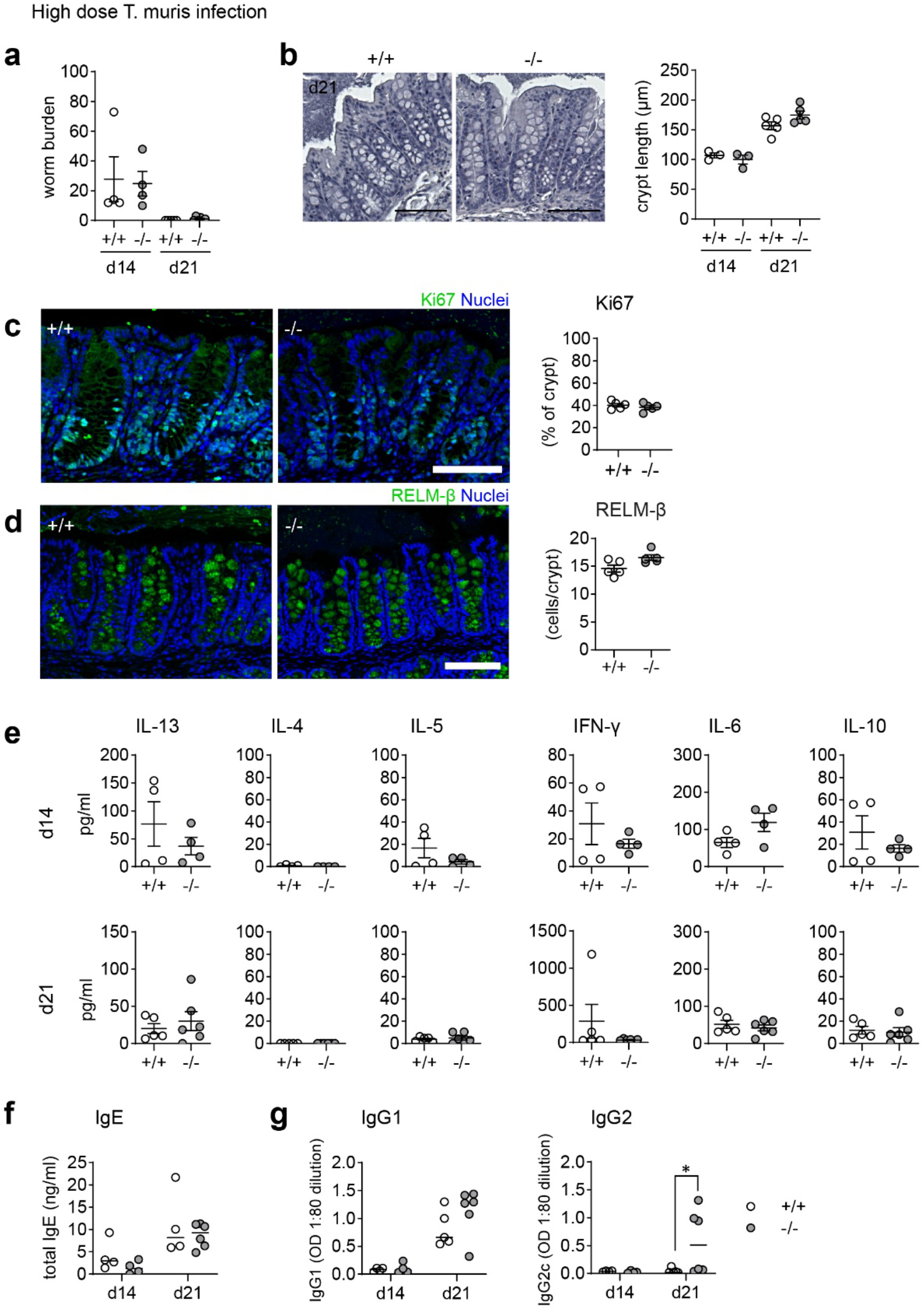
*Mmp17* KO mice show no differences in clearance of high dose *T. muris* infection. (**a**) Graph shows worm burdens in WT (*+/+*) and KO (*-/-*) mice infected with high dose *Trichuris muris* at 14 dpi (d14, 1 experiment n=4) and 21 dpi (d21, 1 experiment,, +/+ (n=5), -/- (n=6)). (**b**) H&E stain of WT (*+/+*) and KO (*-/-*) mouse proximal colon (left) and crypt length measurement in µm (right), of naive and *T. muris* infected (21 dpi) mice naive (n=3), +/+ (n=5), -/- (n=5), scale 100µm. (**c**) Ki67 IF stain of *T. muris* infected WT (*+/+*) and KO (*-/-*) mouse proximal colon 21 dpi, scale 100 µm, Ki67 (green), nuclear stain (blue) on the left, Measurement of Ki67 positive area of the crypt (right). (**d**) Cytokine levels measured via cytokine bead array of mLNLs restimulations with E/S antigen at 14 dpi (n=4) and 21 dpi (+/+ (n=5), -/- (n=6)) for the Th1 cytokines IFNg, IL-6, IL-10, and Th2 cytokines IL-4, IL-5 and IL-13 in pg/ml. (**e**) Graphs show total serum levels of IgE and (**g**) serum levels of parasite specific IgG1 and IgG2c antibodies at 14 (n=4) and 21 (+/+ (n=5), -/- (n=6)) dpi, WT (*+/+*) mice (empty column and circles), KO (-/-) mice (grey columns and circles). Numerical data are means ± SEM. Data was analyzed using Student’s unpaired t-test (two-tailed) (a,c,d,e) and 2way ANOVA (b,e,f,g).

### *Mmp17* KO mice are resistant to low-dose *T. muris* infection without inducing a type 2 immune response

Low-dose *T. muris* infection in C57BL/6 mice leads to a chronic infection due to the induction of a non-protective type 1 immune response.(41) We found that *T. muris*-infected *Mmp17* KO mice are resistant to chronic infection. Compared to WT mice, KO mice exhibited significantly reduced worm burdens (Figure 5a) with 46% (6 out of 13 mice) completely clearing the worms after 35 days, while all WT mice remained infected. At the endpoint (d35), we observed no morphological differences. Measurement of crypt length and Ki67 positive percentage of the crypt revealed no changes in epithelial proliferation (Figure 5b,c). Although we observed increased levels of RELM-β in the proximal colon in the naïve state (Fig 1a,b), we do not see any differences in RELM-β+ cells in the proximal colon of infected mice when comparing WT and KO (Figure 5d). Interestingly, this appears to be the case for all infection models we tested. While we observed significant differences in goblet cell effector expression in healthy KO animals, these differences disappeared during infections as they are strongly induced by the immune response (Figure 4d, Figure 5d).

**Figure 5:**
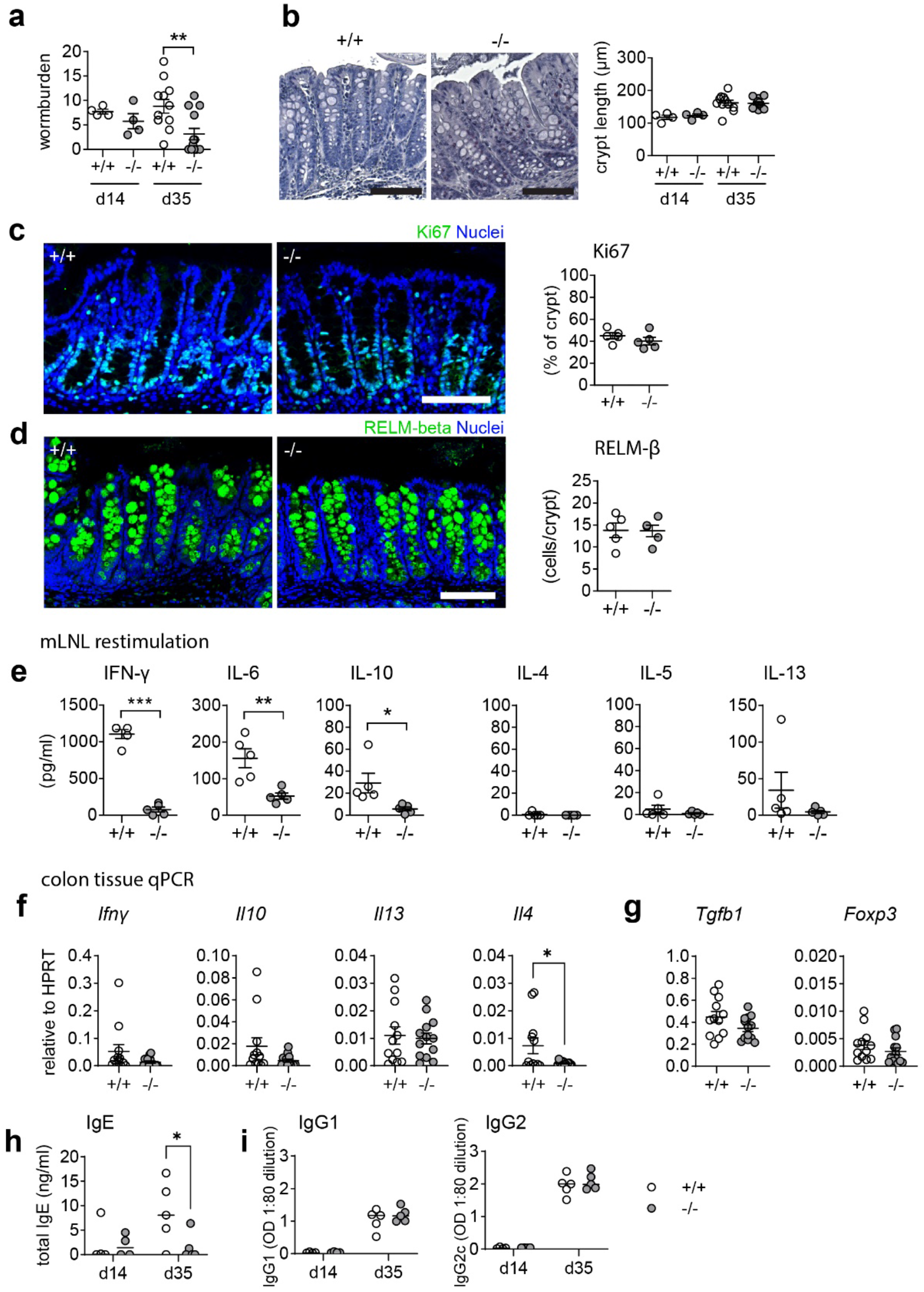
*Mmp17* KO mice exhibit improved clearance in low dose *T. muris* infection. (**a**) The left graph shows the worm burden in WT (*+/+*) and *Mmp17* KO (*-/-*) mice infected with a low dose of *T. muris* at 14 dpi (d14, 1 experiment n=4) and 35 dpi (d35, 2 independent experiments pooled,, +/+: n=12, -/-: n=13). The right graph shows the percentage of mice that cleared the infection/remained infected at 35 dpi. (**b**) Image of an H&E staining of WT (*+/+*) and *Mmp17* KO (*-/-*) mouse proximal colon (left) and crypt length measurement in µm (right), of *T. muris* infected mice at 14 dpi (n=4) and 35 dpi (+/+ (n=12), -/- (n=13)), scale 100 µm. (**c**) IF image of a representative Ki67 stain of *T. muris* infected WT (*+/+*) and KO (*-/-*) mouse proximal colon 35 dpi, scale 100 µm, Ki67 (green), nuclear stain (blue) on the left, Graph of measurements of Ki67 positive part of the crypt (right), n=5. (**d**) RELM-beta IF stain of *T. muris* infected Mmp17 +/+ and -/- mouse proximal colon 35 dpi, scale 100 µm, RELM-beta (green), nuclear stain (blue) on the left, count of RELM-beta positive cells per crypt (right), n=5. (**e**) Cytokine bead array of mLNLs restimulations at 35 dpi (n=5) for the Th1 cytokines IFN-γ, IL-6, IL-10, and Th2 cytokines IL-4, IL-5 and IL-13 in pg/ml, representative data of 1 experiment. (**f**) qPCR of proximal colon tissue at 35 dpi for *Ifng, Il10, Il13*, and *Il4* relative to HPRT, +/+ (n=12), -/- (n=13), pooled from two independent experiments. (**g**) Graphs show mRNA expression data measured using qPCR of proximal colon tissue at 35 dpi for *Tgfb1* and *Foxp3* relative to HPRT, +/+ (n=12), -/- (n=13), pooled from two independent experiments. (**h**) The graphs show total levels of serum IgE antibodies at 14 (n=4) and 35 (n=5) dpi, WT (*+/+*) mice (empty circles), KO (*-/-*) mice (grey circles). (**i**) Graphs depict serum levels of parasite specific IgG1 and IgG2c antibodies at 14 (n=4) and 35 (n=5) dpi, WT (*+/+*) mice (empty circles), KO (*-/-*) mice (grey circles). Numerical data are means ± SEM. Data was analyzed using Student’s unpaired t-test (two-tailed) (a,c,d,e,f,g) and 2way ANOVA (b,h,i).

Infected WT mice exhibited a type 1 cytokine profile consistent with a chronic infection, showing significantly increased levels of IFN-γ, in addition to IL-6 and IL-10, while cytokine levels in KO animals remained low (Figure 5e). It is generally assumed that a type 2 immune response is essential for clearing *T. muris*; however, we did not observe increased levels of type 2 cytokines like IL-4, IL-5 or IL-13 in the KO (Figure 5e) at the time of sampling. To confirm the lack of a local type 2 response in the KO colon, infected colon tissue samples were analyzed for *Ifng, Il10, Il13*, and *Il4* mRNA levels (Figure 5f). Furthermore, tissue mRNA levels of *Tgfb1* and *Foxp3* suggested no involvement of regulatory T-cells in the downregulation of type 1 cytokines in the KO (Figure 5g). Whilst the lack of a detectable Th2 immune response in the KO may reflect the sampling point, and thus a consequence of lower worm burdens at the end point, the significantly lower total serum IgE compared to WT (Figure 5h) suggests that the enhanced resistance to infection seen in the KO mouse is not accompanied by an elevated Th2 immunity. Alcian blue/PAS staining of paraffin sections from infected colon tissue revealed no difference in mucus pH, hinting towards similar mucus quality between WT and KO mice (Supplemental Figure 2a,b). This data collectively demonstrates that *Mmp17* KO mice can clear low-dose *T. muris* infection without mounting a type 2 immune response which is typically seen as requisite for parasite clearance. Thus, we propose that the changes in the naïve state contribute to the enhanced ability of the KO to clear the infection.

### *Mmp17* KO mice show increased goblet cell maturation in cecum crypts

*T. muris* primarily colonizes the cecum and proximal colon(36, 42), therefore, we analyzed if there were also changes in goblet cell effectors in the cecum at steady state. Consequently, we examined the cecum of KO mice, assessing RELM-β and CLCA1 levels and analyzing goblet cell numbers and morphology. Consistent with our findings in the proximal colon (Figure 1d), we detected no differences in goblet cell numbers when comparing WT and KO cecum (Figure 6a,b). However, we observed an increase in goblet cell size in the upper part of the crypt (Figure 6c), suggesting enhanced goblet cell maturation. Upon evaluating goblet cell effectors, both RELM-β and CLCA1 levels were markedly increased in the KO cecum, as demonstrated by IF staining (Figure 6d,e). Moreover, while RELM-β+ cells were predominantly located towards the base of the cecum crypts, CLCA1 signal was mainly present in the upper crypt in WT animals, suggesting that CLCA1 is produced by mature goblet cells.(7) In contrast, in KO mouse cecum, we observed significantly more CLCA1 positive cells toward the lower half of the crypts (Figure 6e).

**Figure 6:**
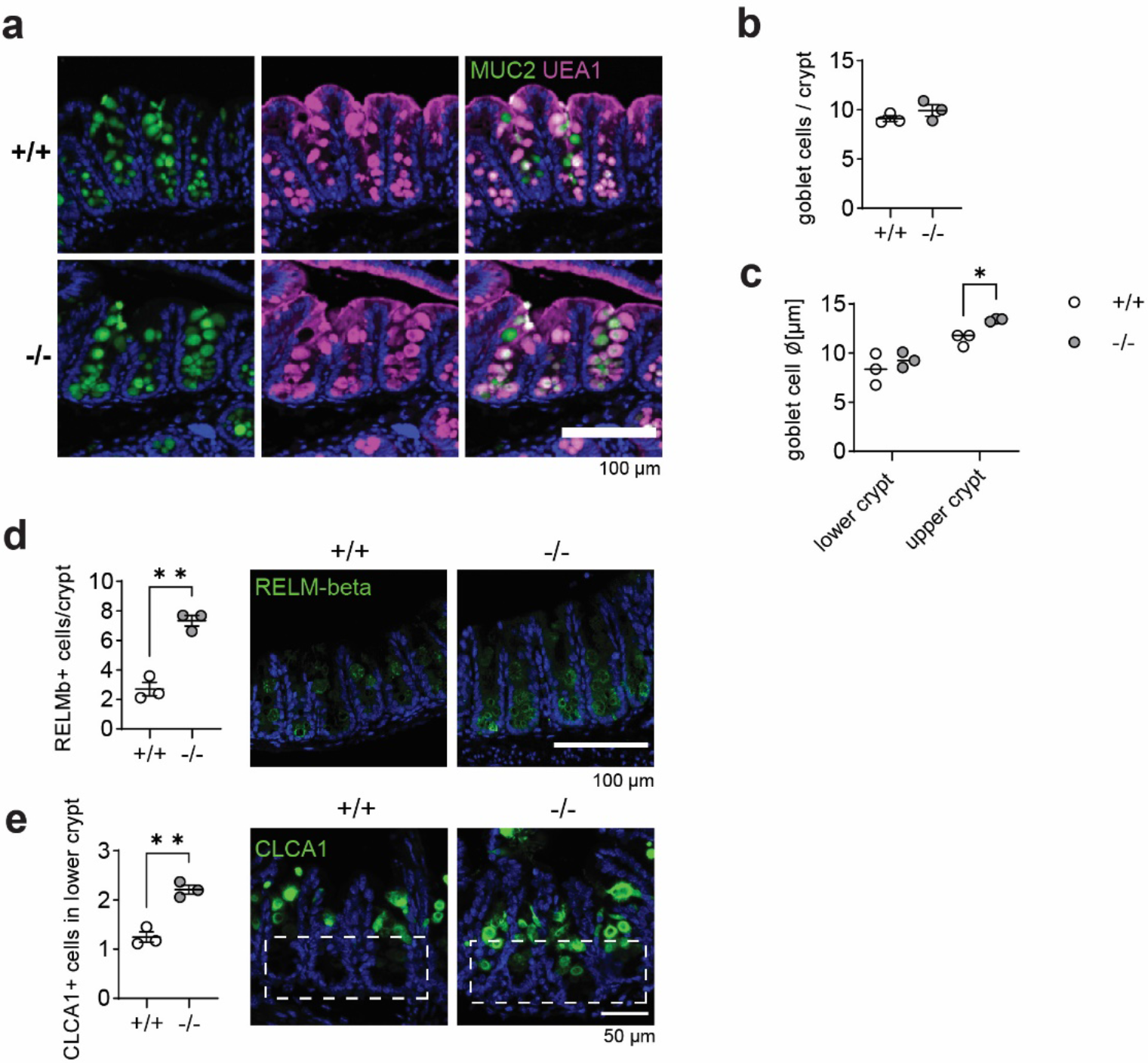
Goblet cells in *Mmp17* KO mouse cecum show increased maturation. (**a**) Muc2 IF and UEA-1 stain of naive mouse cecum of WT (*+/+*) and KO (*-/-*) mice, Muc2 (green), UEA-1 (magenta), nuclear stain (blue), scale bar 100 µm. (**b**) Count of goblet cells per cecum crypts of naive WT (*+/+*) and KO (*-/-*) mice, n=3. (**c**) The graph shows the measurement of goblet cell diameter in lower and upper cecum crypts of naive WT (*+/+*) and KO (*-/-*) mice, n=3, 40-50 cells measured. (**d**) RELM-beta IF stain of naive mouse cecum of WT (*+/+*) and KO (*-/-*) mice, RELM-beta (green), nuclear stain (blue), scale bar 100 µm (right) and count of RELM-beta positive cells per crypt, n=3, min 32 crypts counted (left). (**e**) CLCA1 IF stain of naive mouse cecum of WT (*+/+*) and KO (*-/-*) mice, CLCA1 (green), nuclear stain (blue), scale bar 100 µm (right) and count of CLCA1 positive cells in the lower crypt, n=3, 35 crypts counted (left). Numerical data are means ± SEM. Data was analyzed using Student’s unpaired t-test (two-tailed) (b,d,e) and 2way ANOVA (c).

In accordance with Figure 1, our results suggest an increased goblet cell maturation state in KO mice, which did not correlate with elevated cytokines levels (Figure 1). The change in the zonation of goblet cell effector expression could imply that an altered gradient controlling goblet cell maturation might be the cause for our observations. It has been demonstrated that BMP signaling is in part responsible for goblet cell maturation along the crypt villus axis.(43)

### *Mmp17* is expressed in putative BMP-niche providing cells, and *Mmp17* KO mice have heightened epithelial SMAD4 levels

We have previously reported that the primary cell type expressing *Mmp17* in the colon at steady state are smooth muscle cells, particularly those in the muscularis mucosa, a thin layer of muscle located right beneath the epithelium, which exhibit high levels of *Mmp17* expression.(28) Using the reporter gene β-Galactosidase in the *Mmp17* KO mouse model, we observed lower, yet distinct expression also in non-epithelial cells located in the upper part of the crypt, that do not express the smooth muscle marker alpha-smooth muscle actin (Figure 7a,b). Others have shown that *Mmp17* can also be expressed by monocytes and macrophages in the vasculature. (44) To investigate whether gut macrophages express *Mmp17,* we co-stained with the macrophage marker CD68; however, we did not find co-localization of β-Galactosidase and CD68 (Figure 7a). Instead, we discovered that cells with low β-Galactosidase expression were positive for the mesenchymal marker PDGFRα (Figure 7b). These mesenchymal cells in the intestine play a crucial role in forming the intestinal niche, for instance, by maintaining the BMP gradient through the expression of BMP agonists and antagonists.(16) In a recent paper, Beumer et al. described how the BMP gradient along the crypt-villus-axis influences goblet cell gene expression in the small intestine using both a KO-mouse model and small intestinal organoids.(43) As KO mice have increased goblet cell maturation (Figure 6), but no changes in NOTCH signaling (Figure 1), we were prompted to see if there was altered BMP signaling in the epithelium. Indeed, in KO mice we found an increase in SMAD4 signal in the colon epithelium as an indicator for increased BMP signaling (Figure 7c). Together, we propose that increased BMP signaling is resulting in enhanced goblet cell maturation in *Mmp17* KO mice.

**Figure 7:**
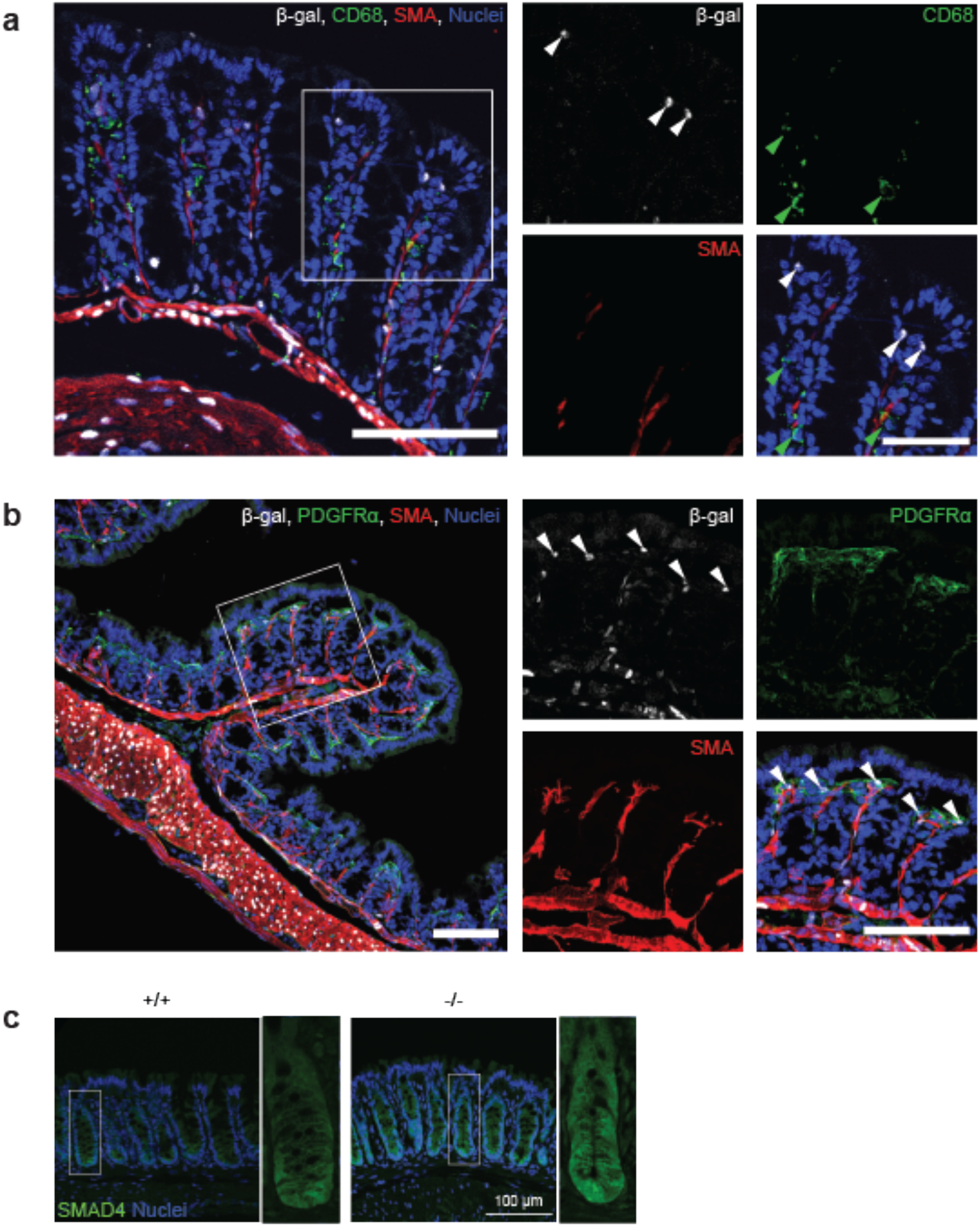
*Mmp17* is expressed in PDGFRα+ mesenchymal cells and *Mmp17* KO mice show elevated BMP signaling in colon crypts. (**a**) β-galactosidase in naive *Mmp17* KO mouse proximal colon, β-gal (white), CD68 (green), smooth muscle actin (SMA) (red), nuclear stain (blue), white arrows point at β-gal+ cells at the tip of the colon crypts. Green arrows point at CD68+ cells, scale bar 100µm, scale bar close up 50µm. (**b**) β-gal in naive *Mmp17* KO mouse proximal colon, beta-Gal (white), PDGFRα (green), smooth muscle actin (red), nuclear stain (blue), arrows point at beta-Gal positive cells at the tip of the colon crypts, scale bars 100µm. (**c**) SMAD4 IF stain of naive proximal colon of Mmp17 WT and KO mice, SMAD4 (green), nuclear stain (blue), scale bar 100 µm.

## DISCUSSION

In the *Mmp17* KO mouse model, we observed an elevated expression of goblet cell effector genes, such as *Ang4*, *Clca1,* and *Retnlb,* in cecum and proximal colon (Figures 1 and 5). While previous studies have shown that mice lacking RELM-β are highly susceptible to *C. rodentium* infection, to our knowledge there have been no investigations into the effect of increased RELM-β levels. Our study did not find any evidence that suggest that the elevated homeostatic levels of MUC2, ANG4, CLCA1, and RELM-β are beneficial during *C. rodentium* infection, as WT and KO mice cleared the infection at the same time, and peak infection numbers were comparable. In a previous study, we reported that *Mmp17* KO mice were highly susceptible to DSS-induced colitis.(28) However, in the case of *C. rodentium* infection, which induces only mild colitis in C57BL/6 mice, we observed no elevated tissue damage in KO mice. This may be due to less overall damage caused by *C. rodentium*-induced colitis, and thus the deficiencies in repair might not be significant in this model. Moreover, both WT and KO mice showed crypt hyperplasia with a comparable average crypt length, indicating a similar proliferation rate.

Resistance to intestinal helminths has been mainly attributed to the type of immune response elicited, with a type 2 response typically leading to clearance, while a type 1 response results in chronic infection.(36) However, our findings suggest that the ability to clear the infection may also be influenced by other factors. *Mmp17* KO mice, infected with a low dose of *T. muris* eggs, were able to clear the infection, while WT C57BL/6 mice developed a chronic infection. Thus, in the KO mouse, worm counts were significantly lower, and almost half of the mice completely cleared the worms by day 35 post infection. Additionally, the KO mice did not show the typical increase in cytokines of a chronic (type 1) infection (IFNg, IL-6, IL-10), nor did they have increased levels of IL-4, IL-13, or IL-5, which would be indicative of a type 2 immune response. Whilst the relative absence of Th2 cytokines, yet enhanced resistance seen in the KO might reflect the sampling point, low cytokine levels were accompanied by subdued total IgE levels in serum which would normally accumulate over time post infection. Further, no differences in epithelial or immune responses were observed in the infected mice. Thus, enhanced resistance is not accompanied by an elevated Th2 immune response. Interestingly a similar disconnect between resistance and Th2 immunity has been reported before in the context of goblet cell derived mucins, with the MUC5ac deficient mice failing to expel *T. muris* despite mounting a strong Th2 response.(45) This could indicate that the changes in the epithelium of KO mice at steady-state may be responsible for the improved clearance of *T. muris* infection in KO mice.

Notably, altered expression of genes involved in mucus production and processing, as well as AMPs were observed in the cecum and proximal colon of KO mice, which are the areas of the intestine that are colonized by *T. muris*. Specifically, there was an increase in goblet cell effectors such as RELM-β and ANG4, both upregulated during helminth infection and RELM-β being able to directly affect the viability of *T. muris* and other parasitic helminths.(14, 46) Additionally, CLCA1, which plays an important role in mucus processing and helps form the two distinct inner and outer mucus layers of the colon by processing mucin 2 (MUC2), is also described to be upregulated during *T. muris* infection.(39) While CLCA1 is not essential in the steady state to form a functional mucus barrier, it is crucial during nematode infections.(47)

One possible mechanism for the improved clearance of *T. muris* infection in KO mice is a thicker mucus layer, which could prevent some larvae invading the epithelium. However, worm burdens at day 14 post infection were equivalent in WT and KO evidencing equivalent establishment. Thus the significantly lower worms burdens seen at day 35 implicate mucus as an aid in flushing worms out in concert with other clearance mechanisms like increased epithelial proliferation and muscle contractions.(48) The increase in goblet cell effectors could also contribute to the enhanced clearance of *T. muris* in KO mice by exerting direct effects on the worms themselves, lowering their viability and hindering their movement. It is of note that while we observe differences in goblet cell effectors in the steady-state, those differences disappear after infection.

Mucus and goblet cell effector genes, such as *Ang4*, *Clca1, Retnlb*, and *Muc2* are typically induced during infections by cytokines, especially IL-4, IL-13 and IL-22. However, we see no indication of increased cytokine levels in the naïve colon or re-stimulated mesenteric lymph node cells that could cause this change (Figure 1g). Furthermore, we did not observe any signs of goblet cell hyperplasia, which is often associated with cytokine-mediated activation (Figures 1c and 5). However, mature goblet cells in the upper part of the crypt appear to be larger in KO animals compared to WT animals, indicating an increase in total mucus production. This finding is consistent with the increased expression of *Muc2* mRNA in KO crypts (Figure 1a). We did not detect any changes in the pH of the mucus, indicating no significant differences in mucus quality (Supplementary Figure 2a,b).

As we could not find detectable differences in cytokine levels in the *Mmp17-/-* colon to explain this changed goblet cell expression pattern there must be another mode of induction. The colon epithelium is regulated by growth factor gradients that help preserve the stem cells at the bottom of intestinal crypts while allowing for mature cells with different functions in the upper part of the crypt. One of those gradients is BMP signaling, kept low by BMP-antagonists at the bottom of the crypt and increasing towards the top. In the *Mmp17* KO mouse colon epithelium we find an increase in BMP-signaling. Although initially thought off as primarily affecting enterocyte maturation, Beumer *et al.* show that BMP signaling can also induce changes in goblet cell expression. Another indication of an altered gradient is the fact we find more CLCA1+ cells in the lower cecum crypts. CLCA1 is a marker for upper crypt goblet cells that are among the main mucus producers in the colon. The presence of CLCA1 towards the bottom of the crypt may indicated a more mature goblet cell state.

Overall, our findings suggest that the altered steady-state with increased expression of goblet cell effectors in the *Mmp17* KO mouse epithelium could be responsible for its ability to clear low dose *T. muris* infection. We find evidence of increased BMP signaling in the intestinal niche that we suspect to be responsible for a more mature and activated goblet cell state. This study contributes to highlighting the importance of MMPs and mesenchymal cells as important regulators of the intestinal niche.

## ACKNOWLEDGEMENTS

We thank the imaging (CMIC) and animal care (CoMed) core facilities at NTNU, as well as the histology, flow cytometry and biological services core facilities at The University of Manchester for assisting in this work. This work was financially supported by the Norwegian Research Council (Centre of Excellence grant 223255/F50, and ‘Young Research Talent’ 274760 to M.J.O. and 326209 to M.M.A) and the Norwegian Cancer Society (182767 to M.J.O. and 245170 to M.M.A.). The Histology Facility equipment in Manchester used in this study was purchased with grants from the University of Manchester Strategic Fund. This work was supported by an MRC grant (MR/N022661/1) awarded to K.J.E. The authors declare that they have no competing interests.

## AUTHOR CONTRIBUTIONS

PMV, MJO, and MMA contributed to conception and design of the study. HTL performed the analyzed the RNAseq data. PMV and MJO wrote the first draft of the manuscript. RF and KJE planned, performed, and analyzed the *Trichuris muris* mouse infections and provided valuable input to the project. PMV, MMA, and NP performed the *Citrobacter rodentium* infections. PMV and MMA provided the histology data and their analysis. PMV performed and analyzed organoid experiments, qPCR, and ELISA. RY performed and analyzed the Western blots. All authors contributed to manuscript revision, read, and approved the submitted version.

## METHODS

### Mmp17-/- mice

Mmp17−/− mice in the C57BL/6 background have been described previously.(49) Mice were handled under pathogen-free conditions in accordance with CoMed NTNU institutional guidelines. Experiments were performed following Norwegian legislation on animal protection and were approved by the local governmental animal care committee.

*Citrobacter rodentium* infection studies were approved by Norwegian Food Safety Authority (FOTS ID: 11842) and were performed in accordance with the guidelines and recommendations for the care and use of animals in research.

Trichuris muris infections were performed at the University of Manchester and were approved by the University of Manchester Animal Welfare and Ethical Review Board and performed within the guidelines of the Animals (Scientific Procedures) Act, 1986. Mice were maintained at a temperature of 20-22°C in a 12h light, 12h dark lighting schedule, in sterile, individually ventilated cages with food and water *ad lib.* Mice were culled via exposure to increasing carbon dioxide levels, an approved schedule 1 method as specified within the guidelines of the Animals (Scientific Procedures) Act.

### Citrobacter rodentium infection

*C. rodentium* infection was performed following established protocols.(50) In brief, a GFP-expressing *C. rodentium* strain was cultured in Luria-Bertani (LB) medium supplemented with 30 µg/ml chloramphenicol at 37°C. Male and female mice, aged 6–10 weeks, were infected with a dose of 10^8^-10^9^ CFU via oral gavage in a total volume of 0.1ml sterile PBS. The mice were monitored daily after the oral administration of *C. rodentium* for weight loss and pain associated behaviors such as hunchbacked posture, piloerection, and rectal prolapse. Fresh fecal pellets were collected, weighed and homogenized in sterile PBS using a FastPrep homogenizer from MP Biomedicals. Dilutions of the homogenate were plated on agar plates supplemented with 30 µg/ml chloramphenicol. The plates were incubated at 37°C overnight and the number of colonies was assessed. At the end of each experiment (12 days post infection or until clearance), mice were euthanized and organ samples were collected for histology, RNA isolation and CFU analysis.

### *T. muris* infection

The parasite was maintained, E/S products generated, and eggs prepared as previously described.(51) Mice were infected with 150 infective T. muris eggs in 200ul by oral gavage. At necropsy the caecum and proximal colon were collected, blinded and stored at -20°C. For worm counts, caecum and colon were defrosted, opened longitudinally and washed in water to remove gut contents. The gut mucosa was scraped off using curved forceps to remove epithelia and worms from the gut tissue. Both the gut contents and removed gut mucosa were then examined and *T. muris* worms counted using a dissecting microscope (Leica S8 APO).

### Mesenteric lymph node (mLN) cells re-stimulation and cytokine bead array

mLNL were brought to cell suspension and 5×10^6^ cells/ml were cultured for 48h at 37°C 5% CO_2_ in RPMI 1640 with 4 hr E/S antigen (50µg/ml). Supernatants were harvested and stored at −20°C until they were assayed for cytokines.

Cytokines were analyzed using the Cytometric Bead Array (CBA) Mouse/Rat soluble protein flex set system (BD Bioscience) in accordance with the manufacturer’s instructions. Cell acquisition was performed on a FACS Verse (BD Biosciences) and analyzed utilizing FCAP Array v3.0.1 software (BD Cytometric Bead Array) was used.

### Mesenteric lymph node lymphocyte restimulation and cytokine ELISA

The mesenteric lymph nodes were dissected from naïve or *C. rodentium* infected mice and kept in sterile PBS. The lymph nodes were passed through 70 μm cell strainer to form a single cell suspension. The cell viability was assessed using Trypan blue and a Countess automated cell counter (Invitrogen). A 96 well plate was pre-incubated with a 5 µg/ml solution of anti-mouse CD3e (100314, Biolegend) in sterile PBS at 37°C for 2 h. The antibody solution was then removed and the isolated lymph node cells were added at 2×10^5^ cells per well together with soluable anti-mouse CD28 (102112, Biolegend) at a final concentration of 2 µg/ml. After incubating the cells for 4 days at 37°C, 5% CO_2_, the cell-supernatant was collected for cytokine analysis by ELISA.

Supernatants from restimulated mesenteric lymph node lymphocytes and tissue homogenates were assessed for IL-13(88-7137, Invitrogen), IL-17A (88-7371, Invitrogen), IL-22 88-7422, Invitrogen) and IFNγ (88-7422, Invitrogen) cytokine levels by ELISA. The kits were used following the manufacturers protocol.

### Total IgE ELISA

Serum was assayed for IgE antibody production. 96 well plates were coated with purified anti-mouse IgE (2ug/ml, Biolegend, Clone: RME-1) in 0.05M carbonate/ bicarbonate buffer and incubated overnight at 47C. Following incubation, plates were washed in PBS-Tw and non-specific binding blocked with 3% BSA (Sigma-Aldrich) in PBS for 1 hour at room temperature. Block was removed and diluted serum (1:10) added to the plate and incubated for 2hrs at 377C. After washing HRP conjugated goat anti-mouse IgE (1ug/ml; Bio-rad) was added to the plates for 1 hour. Finally, plates were washed and developed with TMB substrate kit (BD Biosciences, Oxford, UK). The reaction was stopped using 0.18M H2SO4, when sufficient colour had developed. The plates were read by a Versa max microplate reader (Molecular Devices) through SoftMax Pro 6.4.2. software at 450 nm, with reference of 570 nm subtracted.

### IgG ELISA

Serum was analyzed for parasite specific IgG1 and IgG2c antibody production. 96 well plates were coated with 5µg/ml *T. muris* overnight E/S antigen overnight, plates were washed, and non-specific binding blocked with 3% BSA (Sigma-Aldrich) in PBS. Following washing, plates were incubated with serum (2 fold dilutions, 1:20-1:2560) and parasite specific antibody was measured using biotinylated IgG1 (BD Biosciences) or IgG2c (BD Biosciences) antibodies which were detected with SA-POD (Roche). Finally, plates were washed and developed with TMB substrate kit (BD Biosciences, Oxford, UK) according to the manufacturer’s instructions. The reaction was stopped using 0.18 M H_2_SO_4_, when sufficient colour had developed. The plates were read by a Versa max microplate reader (Molecular Devices) through SoftMax Pro 6.4.2. software at 450 nm, with reference of 570 nm subtracted.

### Sample preparation for histology

Proximal colon tissue of *T. muris* infected mice was fixed for 24 hours in 10% neutral buffered formalin (Fisher) containing 0.9% sodium chloride (Sigma-Aldrich), 2% glacial acetic acid (Sigma-Aldrich) and 0.05% alkyltrimethyl-ammonium bromide (Sigma-Aldrich) prior to storage in 70% Ethanol (Fisher) until processing. Fixed tissues were dehydrated through a graded series of ethanol, cleared in xylol and infiltrated with paraffin in a dehydration automat (Leica ASP300 S) using a standard protocol. Specimens were embedded in paraffin (Histocentre2, Shandon). Alternatively, proximal colon tissue was embedded in OCT and snap-frozen on dry ice and stored at -80C until processing.

For naïve and C. rodentium infected samples, the full colon or cecum tips were dissected, washed with ice-cold PBS. The colon was rolled into “swiss rolls” as previously described or a 5 mm piece of the proximal colon was taken.(52) The tissues were fixed in 4% formaldehyde for 24h at RT (Room Temperature). After the fixation, the tissues were embedded into paraffin.

### Histological stainings

Paraffin embedded tissues were cut into 4µm sections and placed on glass slides. The slides were heated at 60°C for 20 min prior to deparaffinization in Neo-Clear Xylene Substitude (sigma-aldrich). Rehydration was performed through a decreasing ethanol gradient.

For Hematoxylin and eosin stain, the samples were incubated for 1 min in hematoxylin solution. The tissues were then washed with water until the water ran clear and incubated in Eosin Y solution for 3 min.

For Alcian blue stain, the slides were incubated in alcian blue solution for 30 min. After thoroughly washing the slides with water, the tissues were counter stained with nuclear fast red solution for 5 minutes.

For Alcian blue/PAS stain, the tissues were stained in 1% alcian blue in 3% acetic acid (pH2.5) for 5 min, 1% periodic acid for 5 min, Schiff’s reagent for 15 min and finally in Mayer’s hematoxylin for 1 minute. After each staining step the slides were washed in distilled water for 5 minutes.

After the staining the tissues were washed, then dehydrated using an alcohol gradient, cleared in Xylene/Neo-Clear and mounted.

### Tissue immunostainings

The paraffin-embedded or frozen OCT-embedded tissues were cut into 4 μm sections and placed on glass slides. Paraffin-embedded tissue sections underwent deparaffinization, rehydration, and antigen retrieval using citrate buffer at pH-6.0. Frozen sections were fixed in −20 °C cold methanol for 10 min before continuing with blocking. Tissues were then blocked and permeabilized with PBS containing 0.2% Triton X-100, 1% BSA, and 2% normal goat serum (NGS) for 1 hr at RT in a humidified chamber.

Primary antibody incubation was performed using a solution containing 0.1% Triton X-100, 0.5% BSA, 0.05 % Tween-20, and 1% NGS in PBS overnight at 4°C. The following anti-mouse primary antibodies were used: anti-RELM-beta (1:500, rabbit polyclonal antibody, Peprotech, 500-p215), anti-MUC2 (1:200, Santa Cruz Biotechnology, sc-15334), anti-CLCA1 (1:200, rabbit monoclonal antibody, Abcam, ab180851), anti-CD68 (1:200, BioRad, MCA1957T), anti-Ki67 (1:200, rabbit monoclonal antibody, Invitrogen, MA5-14520) anti-SMAD4 (for frozen sections, dilution 1:400, Cell signaling, 46535), anti-βGalactosidase (for frozen sections, dilution 1:100, Rabbit polyclonal antibody, Abcam, ab4761), anti-PDGFR-alpha (15 µg/mL, goat polyclonal antibody, R&D systems, AF1062).

Following primary antibody incubation, slides were washed in washing buffer (0.2% Tween-20 in PBS) three times for 10 min each and then incubated with the appropriate Alexa Fluor secondary antibodies, UEA-1 (1:500, Vector Laboratories, RL-1062), anti-SMA-Cy3 antibody (1:500, mouse mAb, Sigma-Aldrich C6198), and counterstained with Hoechst 33342 (1:1000, MERCK, H6024) for 1 hr at RT. After washing the slides three times with washing buffer and once with distilled water, they were mounted in Fluoromount G (Invitrogen, 00-4958-02). Imaging was conducted with a Zeiss Airyscan confocal microscope, using a 10× and 20× objective lens. Images were analyzed using ImageJ software.

### Colon crypt isolation

Colon crypts for RNA sequencing and organoid culture were isolated using a previously published method (REF). Briefly, the entire colon was flushed with cold PBS and then cut into 2-4 mm pieces. These pieces were washed with cold PBS until the supernatant appeared clear and subsequently transferred into cold chelation buffer (PBS containing 2 mM EDTA) for incubation at 4 °C under gentle agitation for 1 h. The chelation buffer was then removed, and the tissue pieces were resuspended in PBS. Crypts were separated from the tissue by vigorously pipetting with a BSA-coated 10 ml pipette or by shaking the tube. The tissue pieces were allowed to settle at the bottom of the tube for approximately 1 min, after which the supernatant was collected into a BSA coated 50 ml tube. The pipetting/shaking process was repeated using fresh PBS until most crypts had detached. The collected supernatant was centrifuged at 200 x g for 3 min, and the resulting crypt pellet was resuspended in basal crypt medium (BCM, see below) and counted. For subsequent experiments, the crypts were either used for RNA isolation, protein extraction or colonoid culture, seeded into 40 µl drops of Matrigel in a P24 well plate (300 crypts/well) and cultured in WNR medium (medium composition is described below) with Rock inhibitor (10 µM) for 24 h at 37 °C and 5% CO_2_.

### Mouse colon organoid culture

Mouse colon organoids (colonoids) were cultured in WNR medium composed of BCM (advanced Dulbecco’s modified Eagle medium-F12 supplemented with penicillin/streptomycin, 10 mM HEPES, 2 mM Glutamax) plus 1× N2 (ThermoFisher Scientific 100×, 17502048), 1× B-27 (ThermoFisher Scientific 50X, 17504044), and 1× N-acetyl-L-cysteine (Sigma, A7250). The medium was overlaid with 50 ng/ml of murine EGF (Thermo Fisher Scientific, PMG8041) and 50% L-WRN (ATCC 3276) cell line conditioned medium, enriched in Wnt, R-sponding, and Noggin.(15) After isolation or splitting the crypts/colonoid pieces were seeded into 40 µl drops of Matrigel in a P24 well plate (300 crypts/well) and cultured in WNR medium (medium composition is described below) with Rock inhibitor (10 µM) for the first 24 h at 37 °C and 5% CO_2_. After that the medium was replaced every second or third day. For passaging, Matrigel and organoids were disrupted mechanically via vigorous pipetting. The suspension was centrifuged at 300 × g, 5 min at 4 °C and the pellet was resuspended in fresh, cold Matrigel to be plated in P24 wells or eight-well ibidi chambers (80821, Ibidi) for immunostainings. For differentiation, the medium was switched to ENR medium consisting of BCM plus 1× N2, 1× B-27, and 1× N-acetyl-L-cysteine supplemented with 50 ng/ml murine EGF, 20% R-Spondin-conditioned medium (kindly rovided by Calvin Kuo, Stanford University School of Medicine, Stanford, CA, USA), and 10% Noggin-conditioned medium after 3-4 days of culture.

### Organoid immunostaining

Organoid immunostaining was performed using a previously described protocol.(15) In brief, organoids were fixed with 4% paraformaldehyde with 2% sucrose for 30 min at room temperature. After washing with sterile PBS, the organoids were permeabilized with 0.2% Triton X 100 in PBS for 30 min at room temperature. Blocking was performed for 1 hour at room temperature in PBS containing 1% BSA, 2% NGS, and 0.2% Triton X 100. Primary antibodies (RELMβ (1:200, Peprotech 500-p215)) were diluted in the same blocking buffer and incubated with the organoids over night at 4 °C. After washing, the organoids were incubated with the appropriate Alexa Fluor secondary antibodies, UEA-1 (1:500, Vector Laboratories, RL-1062), and counterstained with Hoechst 33342 (1:1000, MERCK, H6024) over night at 4 °C in the dark. After incubation, the organoids were washed and 250 µl Fluoromount G was added to each well. Imaging was conducted with a Zeiss Airyscan confocal microscope, using a 10× and 20× objective lens. Images were analyzed using ImageJ software.

### RNA seq of colon crypts

RNAseq data from colon crypts and muscle from Mmp17 KO mice was obtained and analyzed as described in our previous publication.(28) In brief, colon crypts were isolated using the colon crypt isolation protocol above. The crypts were placed in lysis buffer (RLT, provided in RNeasy® Mini Quiagen Kit, 74104) and RNA was isolated following manifacturer instructions (RNeasy® Mini Kit, Qiagen, 74104). RNA quantification was done by spectrophotometry (ND1000 Spectrophotometer, NanoDrop, Thermo Scientific) and 25 µl with an RNA concentration of 50 ng/µl were used for RNA seq. The library preparation and sequencing was performed by the NTNU Genomic Core facility using a Lexogen SENSE mRNA library preparation kit to generate the library and Illumina NS500 flow cells to sequence the samples at 2 x 75 bp paired end. The STAR aligner was used to align reads to the Mus musculus genome build mm10. featureCounts was used to count the number of reads that uniquely aligned to the exon region of each gene in GENCODE annotation M18 of the mouse genome. Genes with less than 10 total counts were filtered out and DESeq2 with default settings was used for differential expression analysis.

All raw sequencing data are available online through ArrayExpress: WT and KO smooth muscle and crypt RNA seq: E-MTAB-9180.

### qPCR

RNA was isolated from homogenized colon tissue (fresh, stored in RNAlater or Trizol) or isolated crypts using the Quick-RNA Miniprep Kit (R1055, Zymo Research), or for samples in Trizol, the Direct-zol RNA Miniprep kit (R2051, Zymo Research), following the manufacturer’s protocols. Colonoid RNA was isolated using the Quick-RNA Microprep Kit (R1050, Zymo Research), following the manufacturer’s protocol. RNA concentration was measured using a spectrophotometer (ND1000 Spectrophotometer, NanoDrop, Thermo Scientific).

The purified RNA was transcribed to cDNA using the Applied Biosystem High-Capacity RNA-to-cDNA Kit. Quantitative real-time PCR was performed with the StepOnePlus Real-Time PCR System using SYBR Green PCR Master Mix by Applied Biosystem. All primers are listed in supplementary table 1.

### Western blot

Pelleted crypts were kept at -80°C and lysed with RIPA buffer supplemented with fresh cOmplete™ protease inhibitor cocktail (Roche 11836170001) and PhosSTOP™ phosphatase inhibitor cocktail (Roche 4906837001). Pellets were then lysed using vortexing and ultrasound sonication, then centrifuged to remove debris. Total protein of the lysate was quantified using BCA assay (Pierce 34335). 100ug of total protein was loaded onto each well of a NuPAGE™ 4 to 12%, Bis-Tris gel (Invitrogen WG1402BOX) and run in MOPS buffer for 20 min at 100 V, then 1 h at 185 V. Transfer from gel to nitrocellulose was done with the Invitrogen iBlot2 Gel Transfer Device with 20 V for 7 min. Blots were blocked with 5% BSA or 5% nonfat dry milk in TBS-T according to manufacturers’ recommendations, for 1h at room temperature, then primary antibodies (Stat3 (Cell Signaling Technology 9139S; 1:1000 in 5% nonfat dry milk and TBS-T), Stat6 (Cell Signaling Technology 5397S; 1:1000 in 5% BSA and TBS-T), pStat3 (Cell Signaling Technology 9145S; 1:2000 in 5% BSA and TBS-T), pStat6 (Cell Signaling Technology 56554S, 1:1000 5% BSA and TBS-T), and GAPDH (Abcam ab125247; 1:10000 in 5% nonfat dry milk and TBS-T)) were added and incubated with gentle shaking overnight at 4°C. On the following day, blots were washed before adding the secondary antibodies (Polyclonal Swine Anti-Rabbit Immunoglobulins/HRP (Dako p0399; 1:5000 in 1% BSA and TBS-T) and Polyclonal Goat Anti-Mouse Immunoglobulins/HRP (Dako p0447; 1:5000 in 1% BSA and TBS-T)) and incubated for 1 h at room temperature with gentle shaking. After washing the blots were developed for 3 minutes using SuperSignal™ West Femto Maximum Sensitivity Substrate (Thermo Scientific 34094). Chemiluminescence was read using a LI-COR Odyssey Fc Imager with 1 min exposure for GAPDH, 2 min exposure for total STAT proteins, and 30 min exposure for phospho-STAT proteins.

## SUPPLEMENTARY MATERIALS

**Supplementary figure 1:**
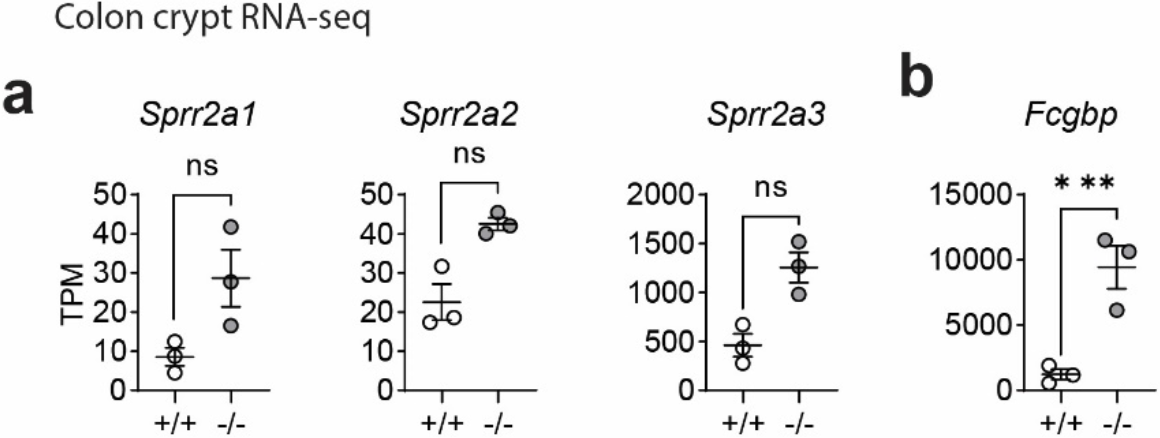
MMP17 controls goblet cell gene expression in Colon. (**a**) RNAseq data of naive mouse colon crypts from WT (*+/+*) and *Mmp*17 KO (*-/-*) mice showing the expression of for *Sprr2a1, Sprr2a2, Sprr2a3* and *Fcgbp* (**b**), n=3, pa dj * p<0.05, ** p<0.01, *** p<0.00. Numerical data are means ± SEM. Data represents p-adjusted value from RNAseq analysis.

**Supplementary Figure 2:**
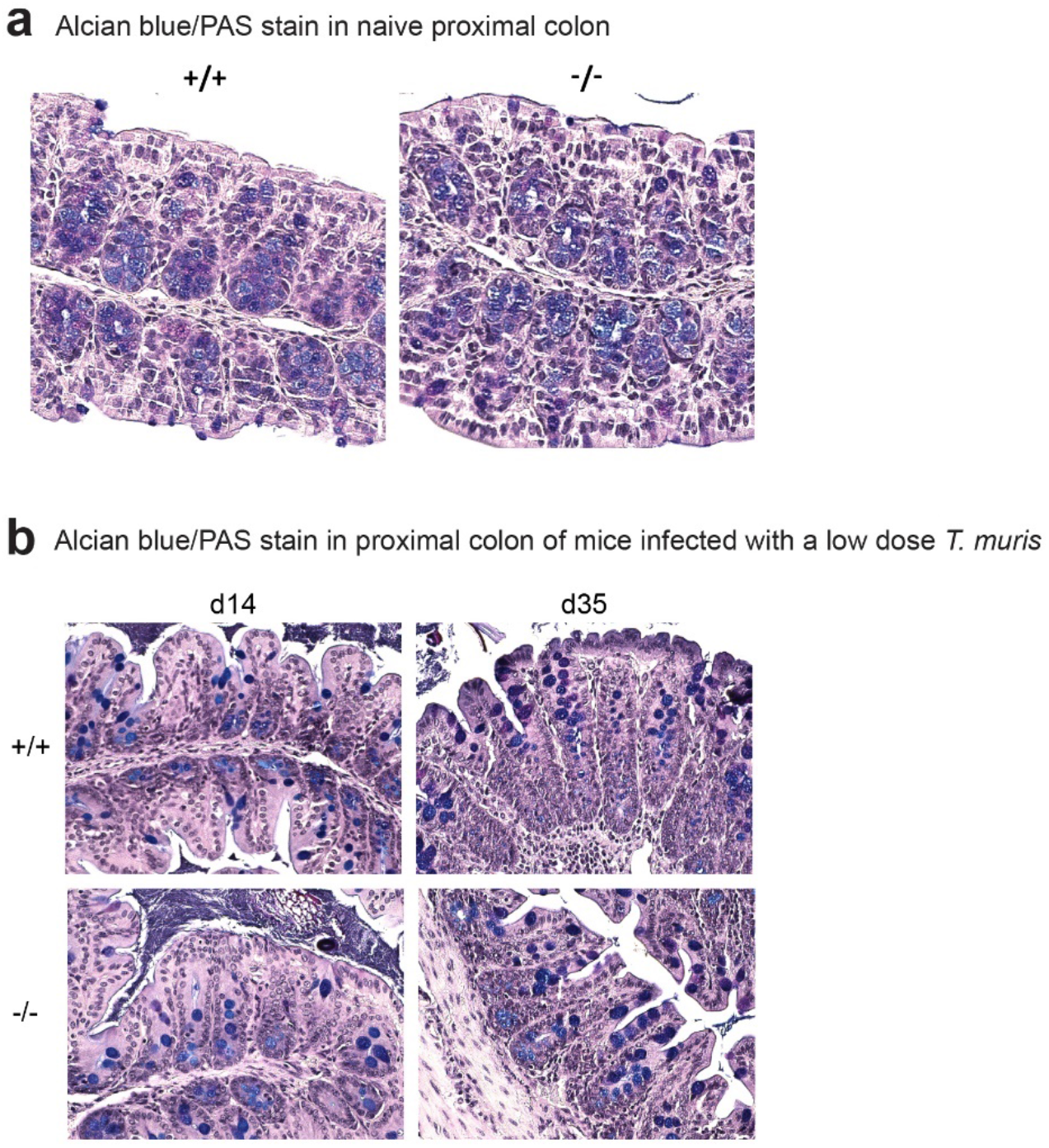
Alcian blue/PAS stain of naive mouse proximal colon of WT (*+/+*) and *Mmp17* KO (*-/-*) mice at 14 dpi and 35 dpi.

**Supplementary table 1:**
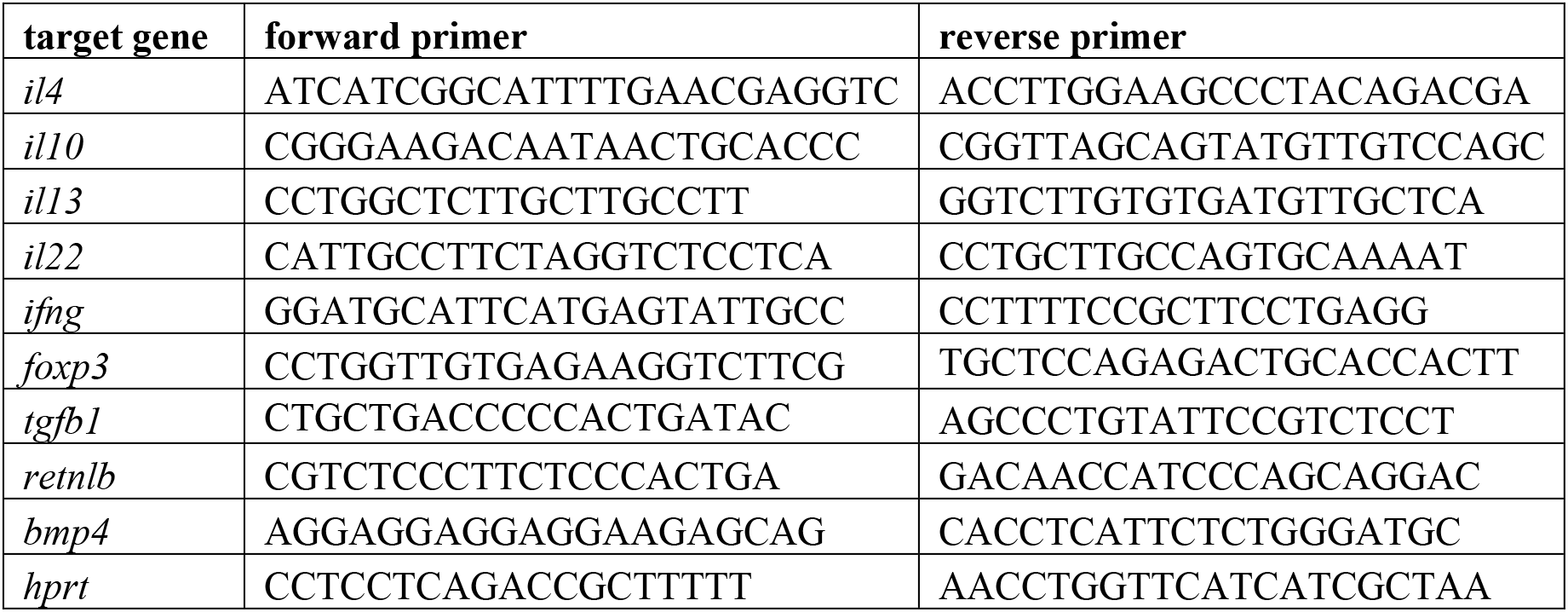
Primers used for qPCR.

